# Optimization of heuristic logic synthesis by iteratively reducing circuit substructures using a database of optimal implementations

**DOI:** 10.1101/2025.11.28.691216

**Authors:** Anh Phong Tran, Dhruv D. Jatkar, M. Ali Al-Radhawi, Elizabeth A. Ernst, Eduardo D. Sontag

**Affiliations:** Department of Chemical Engineering, Northeastern University, Boston, MA 02115 USA; Department of Bioengineering, Northeastern University, Boston, MA 02115 USA; Department of Electrical and Computer Engineering, Northeastern University, Boston, MA 02115 USA; Department of Mathematics, Statistics, and Computer Science, Macalester College, Saint Paul, MN 55105 USA

**Keywords:** Heuristic logic minimizer, Boolean circuit reduction, optimal synthesis, logic optimization, synthetic biology

## Abstract

Minimal synthesis of Boolean functions is an NP-hard problem, and heuristic approaches typically give suboptimal circuits. However, in the emergent field of synthetic biology, genetic logic designs that use even a single additional Boolean gate can render a circuit unimplementable in a cell. This has led to a renewed interest in the field of optimal multilevel Boolean synthesis. For small numbers (1-4) of inputs, an exhaustive search is possible, but this is impractical for large circuits. In this work, we demonstrate that even though it is challenging to build a database of optimal implementations for anything larger than 4-input Boolean functions, a database of 4-input optimal implementations can be used to greatly reduce the number of logical gates required in larger heuristic logic synthesis implementations. The proposed algorithm combines the heuristic results with an optimal implementation database and yields average improvements in fractional gate-count reduction relative to ABC of 5.16% for 5-input circuits and 4.54% for 6-input circuits on outputs provided by the logic synthesis tool *ABC*. In addition to the gains in the efficiency of the implemented circuits, this work also attests to the importance and practicality of the field of optimal synthesis, even if it cannot directly provide results for larger circuits. The focus of this work is on circuits made exclusively of 2-input NOR gates but the presented results are readily applicable to 2-input NAND circuits as well as (2-input) AND/NOT circuits. The framework proposed here is likely to be adaptable to other types of circuits. Moreover, a small computational pipeline is provided for integration with synthetic biology tools such as *Cello*. An implementation of the described algorithm, *HLM* (Hybrid Logic Minimizer), is available at https://github.com/sontaglab/HLM/.

## I. Introduction

The translation of a behavioral model into a circuit layout using automatic synthesis is a central computer-aided design (CAD) problem. The problem of logic synthesis is generally divided into two categories: two-level synthesis and multi-level synthesis. Two-level synthesis is considered a mature field as the optimal implementation of a mapping with a large number of inputs can be computed in a reasonable amount of time [1], [2]. For circuits of small size, the Quine-McCluskey algorithm was standard, but was later replaced by the more efficient Espresso heuristic suboptimal logic minimizer [3]. This is particularly useful when designing programmable logic arrays (PLAs) but typical circuit designs require multiple levels of logic. Due to the larger degrees of freedom that multilevel synthesis algorithms need to deal with, the search for an optimal solution is considerably more challenging, even for a 5-input Boolean function [4]. For these reasons, there has been little interest in recent decades in the area of optimal multi-level synthesis, with most of the contributions having been made in the 1950’s through 1970’s [5]–[15].

In contrast, the heuristic approach to multi-level synthesis has been a major focus of the field and makes a trade-off of the optimality of the solution in favor of other criteria and the ability to find large circuit implementations in a reasonable amount of time. In the 1970’s with the IBM and *LSS* system [16], the rule-based approach was introduced and leveraged local transformations based on ad hoc rules and based on patterns. This had the downside of missing the global view of the synthesis problem. The 1980’s saw the rise of the algorithmic transformations with algorithms such as *MIS* [17] and *BOLD* [1], [18]–[24]. Progressively, these algorithmic transformations were combined with the rule-based approach by using the former in the initial stages of the circuit design and using the latter in the later stages. Approaches based on this hybrid approaches include *SOCRATES* [18] and more recent version of *LSS* [23]. The *SIS* logic synthesis flow was built upon *MIS*, while bringing its own series of improvements [25], [26]. Around the same time as *SIS* in 1990’s, the multi-valued logic synthesis system (*MVSIS*) [27] introduced the more efficient logic representations known as And-Inverter Graphs (AIGs) that later led to the developments of *ABC*, that took the best of *MVSIS* and added new features and improvements [28], [29]. Conceptually, our enumeration of ≤ 4-input cones parallels *k*-cut enumeration and DAG-aware AIG rewriting used in FPGA/LUT flows [29]. Our contribution is complementary: instead of heuristic template-based rewrites, each selected cut is replaced by a functionally equivalent *optimal* (minimum-NOR2) implementation drawn from a precomputed database. The majority of the above works have been developed to cater to the needs of electronic circuit design, focused on scalable synthesis of large circuits. In contrast, we are motivated by the emergent technology of genetic logic design [30], [31], [32], [33]. Despite the vital applications that can be tackled by this nascent field [34], its promise is hampered by severe limitations on the number and the types of gates that can be used inside a biological cell. Although the field has achieved long strides by the adoption of CRISPRi-based repressors to implement NOT and NOR gates [31], [35], [32], [36], it uses a protein known as dCas9 that is toxic to the cell at high concentrations [37]. Therefore, the number of logical 2-input NOR gates per cell has not exceeded 7-8 gates as of now [35], [38]. In this context, a difference of even one gate can determine whether a logical circuit is feasible for genetic implementation or not, which underscores the importance of optimal and near-optimal synthesis.

In this work, we consider the problem of multilevel Boolean synthesis using 2-input NOR gates only. To that end, we introduce an algorithm that takes the best of both the heuristic and optimal multi-level synthesis and combines it into a hybrid framework that yields more efficient circuits. The optimal circuit implementations are based on the thesis work of Ernst [4] and are used to improve the results provided by *ABC* and make the final designs significantly more efficient for medium size circuits (defined here as 5–6 primary-input networks.

## II. Methodology

### A. Database of optimal synthesis implementations

In the context of this work, the definition of “optimal” pertains to the optimality of a circuit (in the sense that it has a minimal number of unique gates among all the circuits implementing a given function). The Branch and Bound Exact Synthesis System (*BESS*) algorithm was used to develop a database of optimal NAND with fan-in of 2 (NAND2) circuits for all 2-, 3-, and 4-input Boolean functions [4], [32]. This database was used to readily generate the corresponding optimal 2-input NOR circuits (NOR2) database using the appropriate circuit transformations. The limitation in building a very large database stems not only from the lengthy exhaustive search that even an individual 5-input Boolean function requires but also from the fact that the number of database entries necessary grows too rapidly to be practical with larger circuits. This is most easily seen by observing that there are 68 P-equivalent 3-input functions, 3,904 4-input functions, and 37,329,264 5-input functions (two Boolean functions are P-equivalent if one function can be transformed into the other by permuting the input variables). In addition to the issue of calculating the individual implementations of Boolean functions, a larger database also causes issues with storage and access. For these reasons, the investigation is focused on using an optimal implementation database for only up to 4-input Boolean functions. Let us note that there can be multiple optimal implementations of the same Boolean logic. The focus of this work is limited to the first implementation that is found through the *BESS* algorithm.

### B. Heuristic logic synthesis

To get the circuit implementations for Boolean functions with more than 4 inputs, the well-established *ABC* tool was used [28]. The commands used in *ABC* in order are: *strash, rewrite, refactor*, and *balance*. Our framework uses *MATLAB* as an interface to create the input files, launch the *ABC* simulations through the command line, to parse the results from the output files, and to convert from an AND-Inverter Graph (AIG) to a NOR-Inverter Graph (NIG).

### C. Hybrid logic minimizer algorithm

The main idea of this algorithm is to look for subcircuits that can be simplified using the optimal synthesis database. A schematic of the algorithm is shown in Fig. 1.

**Fig. 1.**
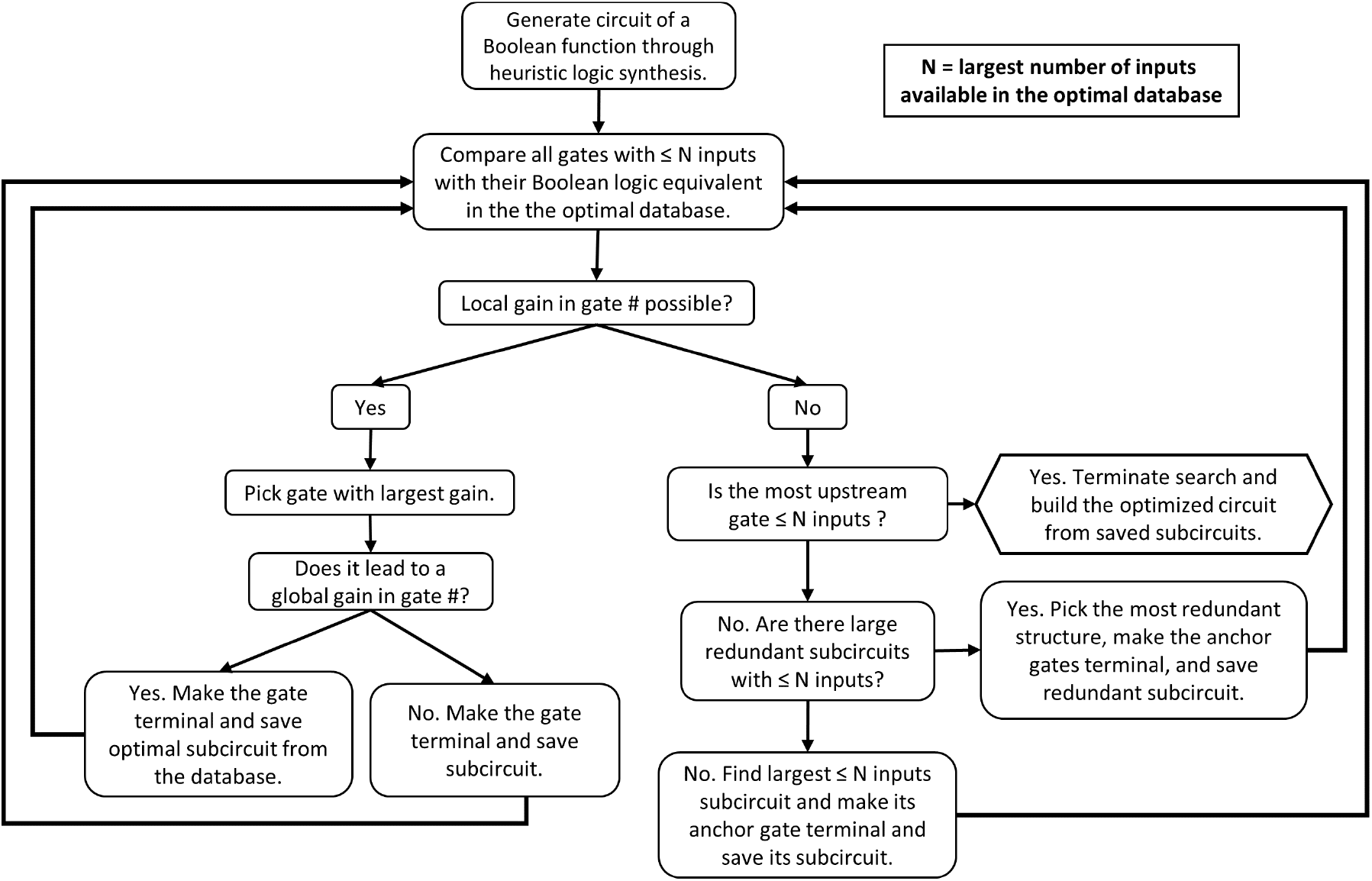
A flowchart of the hybrid logic minimizer algorithm.

Given that the optimal database does not contain entries for circuits containing more than 4 inputs, the focus of this algorithm is limited to substructures of 4 inputs or less. Even with this limited set of substructures, analyzing the possibilities exhaustively would still be problematic. In addition to potentially having to identify all the possible substructures of 4 inputs or less, the order in which these substructures are to be replaced by their optimal counterparts can lead to different outcomes in the observed overall gate reduction. The algorithm presented here does not aim to reduce the number of overall unique gates optimally but rather aims to reduce the search space and finding a “desirable” solution rapidly. The algorithm is subdivided into three steps that are checked successively at each iteration (in an *if, else if, else* pattern):

#### 1) Substitute an eligible substructure with its optimal counterpart

Taking the circuit to be analyzed and thinking of gates in terms of nodes, any parent node that has children nodes totaling more than 4 inputs does not qualify to be optimized at this step. Evidently, any parent node containing a child node that does not qualify also does not qualify. By doing this, the algorithm only has to search substructures that are close to the terminal nodes of the circuit, limiting greatly the complexity of the search. The circuit of each eligible node formed by the node and all of its descendants is then compared with the NOR2 optimal database and the difference of gate numbers is recorded. At the end of this step, the algorithm replaces the circuit with the largest recorded difference in gate numbers by its optimal counterpart. The overall circuit to be considered in subsequent steps has the root of the replaced circuit replaced by a terminal node as the algorithm no longer needs to optimize this part of the circuit. At the next iteration of the search, the size of the considered circuit or netlist is thus reduced.

The optimized circuit is tested at the end of this step by replacing the altered terminal nodes with the corresponding subcircuits to rebuild the entire logic circuit. This ‘global’ implementation of the Boolean logic is used to check whether the local improvement led to a global improvement in terms of reducing the number of unique gates. If it fails to do so, the original substructure that was replaced is saved as a subcircuit instead of the one provided by the database. This ensures that the circuit unique gate count cannot be made worse by this step.

#### 2) Substitute all occurrences of a redundant optimal substructure

When no direct improvement can be found, it also indicates that eligible substructures are found to be optimal with respect to the database. In order to continue shrinking the search space, the algorithm attempts to find substructures to substitute (in other words, replacing the root node of this substructure with a terminal node in the considered circuit). The choice of substructures to be substituted can be quite large. The main issue with replacing a substructure at random is that the root location where the replacement takes place introduces a new input to the overall circuit. It is apparent that having too many unique inputs to a circuit reduces the ability of the algorithm to execute Step 1. Thus, our implementation looks for the 3-input or 4-input substructure that is most redundant in the circuit in terms of occurrences. By doing this, the search space is shrunk, new eligible nodes are created for Step 1, the replaced substructure is optimal, and the same new input is introduced at all the replaced root nodes.

#### 3) Substitute a substructure that maximizes the reduction in the number of unique inputs in the netlist

At this step, there is neither a readily available substructure to optimize nor are there eligible redundant substructures. Let us note that by substituting a substructure that may shrink the number of unique inputs in the considered circuit, it is also increasing the chances that Step 1 finds a substructure to be optimized at the next iteration.

In our implementation, a list of unique inputs is made and the number of times an input appears in the circuit is recorded for each of the unique inputs. This information is then used in the algorithm to remove a 3-input or 4-input substructure that has the lowest mean occurrence across its inputs. In other words, the mean of the input frequency information is used to rank the circuits and remove the circuit with the lowest mean occurrence value. It is no guarantee of reducing the number of unique inputs but it was found to be robust in attempting to remove less frequent inputs. Similarly to Step 2, it also removes an optimal substructure to shrink the overall search space and tries to increase the number of eligible substructures for Step 1.

### D. Overall execution

Steps 1 through 3 are run iteratively until the considered circuit can no longer be shrunk. The complete circuit is then reconstituted from its substituted subcircuits. The details of the implementation are included with the computational package.

A representative example of this algorithm is shown in Fig. 2. While experimentally we are still limited to 7-8 gates in a single cell, circuits up to 110 gates remain biologically relevant for multi-cell systems [39].

**Fig. 2.**
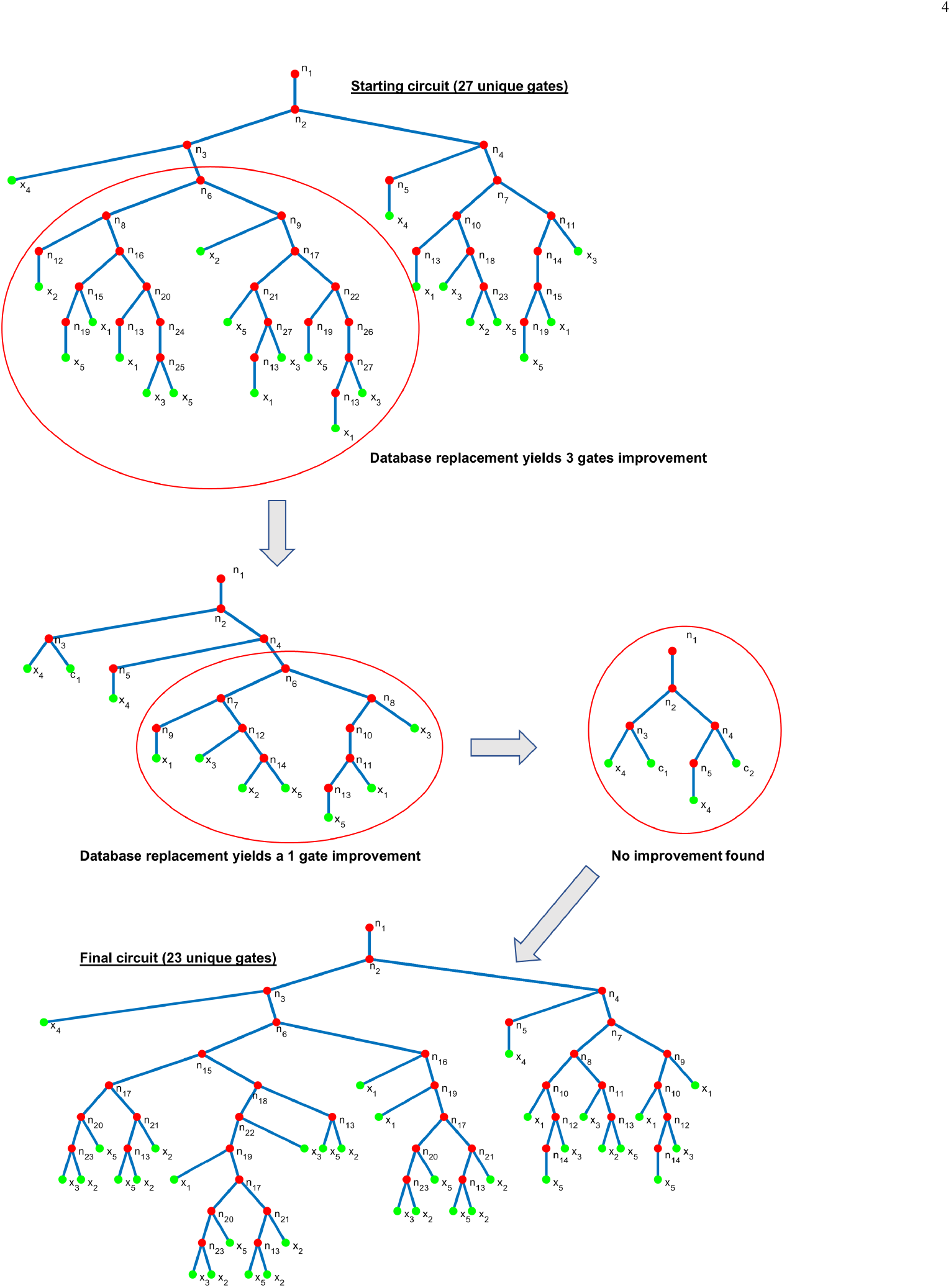
Representative example for the 5-input Boolean function “98546B83” (expressed in hexadecimal). This example shows how the proposed algorithm is able to find an improvement of 4 unique gates. The red circles indicate the current target for improvement by comparison with the optimal database. Each red dot represents a NOR gate and each green dot represents an input or subcircuit *c*_*i*_.

## III. Experimental Results

Before considering the performance of the proposed algorithm, *ABC* was used to generate all the possible 65,536 4-input Boolean functions and the results were compared with the entries of the developed optimal database of NOR2 circuits. These results are summarized in Fig. 3. On average, *ABC* produces circuits that require 4.310 additional gates and only finds the optimal solution 5.630% of the time. The mean number of gates are 9.514 and 13.824 gates for the circuits derived from the optimal synthesis and *ABC*, respectively. From the box plot in Fig 3(d), the trend is clear that larger optimal circuits also lead to a larger deviation from optimality for the *ABC* implementations, primarily due to increased complexity. This gives an idea of what is the theoretical maximum gain in terms of gate number.

**Fig. 3.**
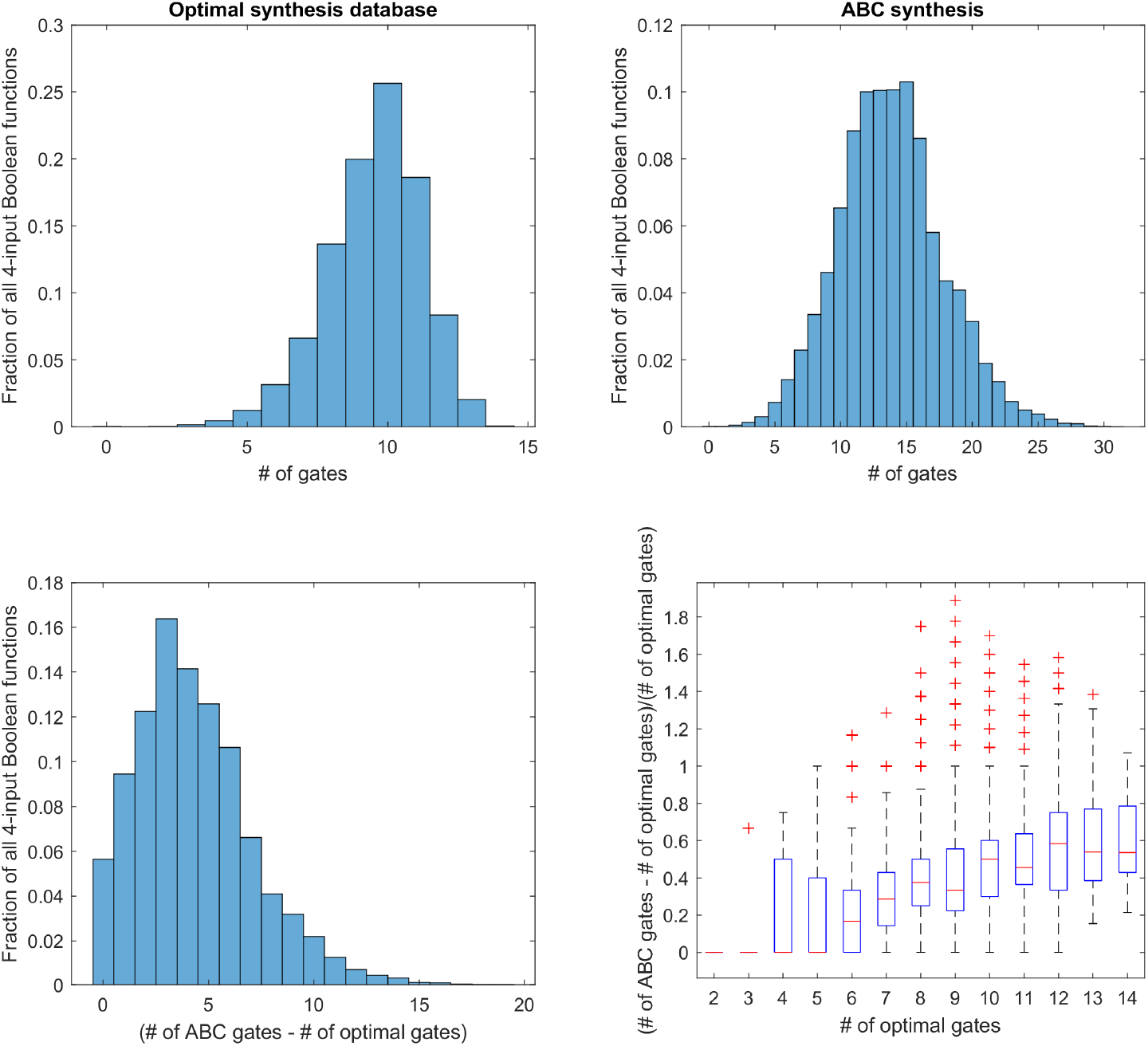
Comparison in the number of gates for NOR2 circuits generated from ABC and those generated through optimal synthesis for all 4-input Boolean functions. (a) and (b) show the distributions in the number of gates for the optimal synthesis database and the ABC synthesis, respectively. (c) shows the distribution in the difference in gate numbers between the two results. In (d), a box plot is drawn of the deviation from optimal normalized by the number of optimal gates as a function of the number of optimal gates.

The algorithm is then tested on 20,000 random 5-input Boolean functions. The average number of gates in a 5-input circuit generated by ABC is estimated to be 28.89 gates. The algorithm yields on average a gain of 1.46 gates. When individually normalizing the gain by the number of gates obtained by ABC, the gain was on average 5.16%. The algorithm did not improve the underlying circuit in 41.68% of cases. Looking only at the improved circuits, the improvement was on average 2.51 gates for an average normalized improvement of 8.85%. These results are also reported as histograms in Fig. 4.

**Fig. 4.**
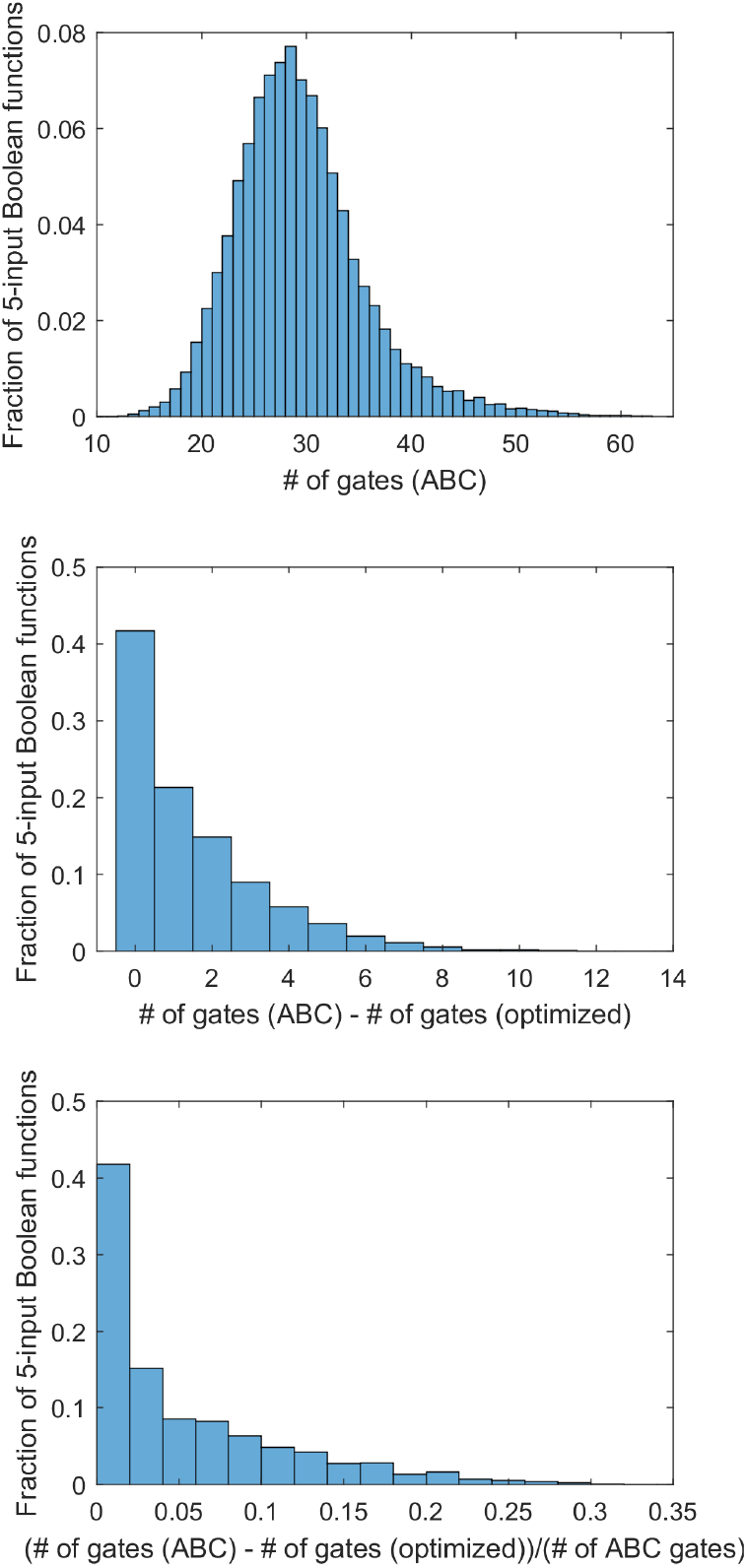
20,000 5-input NOR2 circuits are generated at random from ABC and optimized using the described workflow. (a) shows the distribution of the number of gates in the ABC circuits. (b) is a histogram of the improvements made by the proposed algorithm in terms of the number of gates. (c) is a histogram of the fractional improvement made by the proposed algorithm.

Similarly, the results of the algorithm on 12,000 random 6-input Boolean functions are shown in Fig. 5. The average number of gates for 6-input Boolean functions was found to be about 84.24 gates in *ABC*. The average gain from the algorithm is 3.87 gates or 4.54%. The algorithm fails to improve the *ABC* results in only 17.78% of cases. Looking only at the improved circuits, the gains were 4.71 gates or 5.52%.

**Fig. 5.**
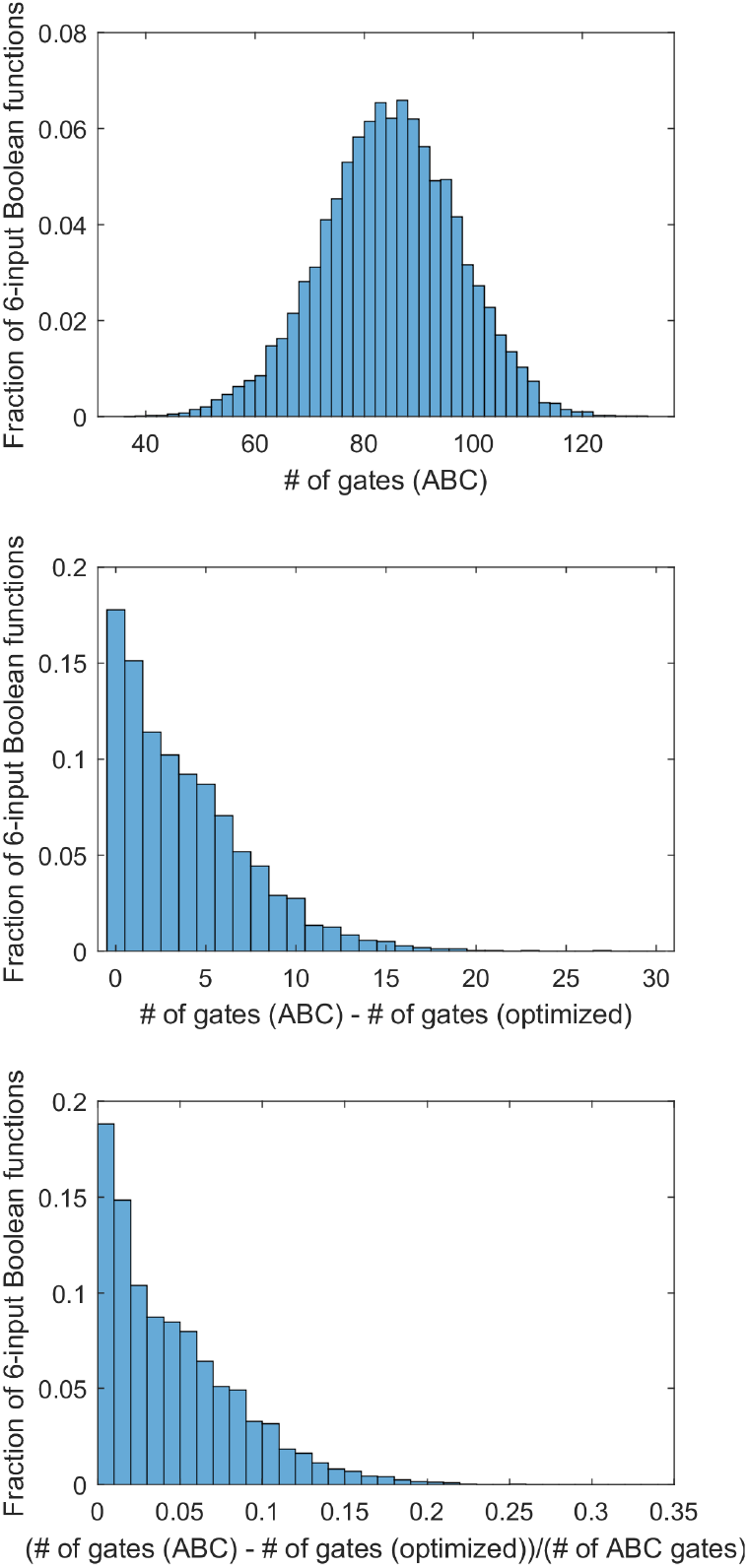
12,000 6-input NOR2 circuits are generated at random from ABC and optimized using the described workflow. (a) shows the distribution of the number of gates in the ABC circuits. (b) is a histogram of the improvements made by the proposed algorithm in terms of number of gates. (c) is a histogram of the fractional improvement made by the proposed algorithm.

**Fig. 6.**
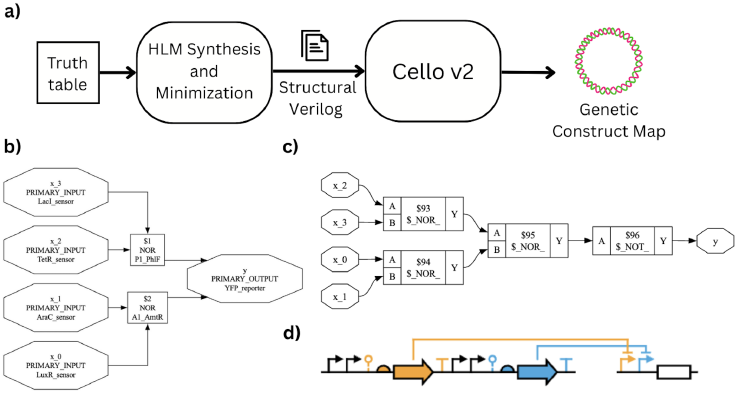
In numerical experiments, the hybrid logic minimizer algorithm performs like a light-weight version of Cello-v2’s built-in network minimizer. (a) gives a high-level description of the computational pipeline. (b) shows the realized circuit with technology mapping (signaling molecules assigned) for the function 0xF888. (c) shows the circuit diagram before being passed to technology mapping. (d) shows the final genetic construct diagram output.

These improvements of 5.16% for 5-input and 4.54% for 6-input Boolean functions are significant given the maturity of the field of logic synthesis. What makes this finding interesting is that a 4-input database requires little memory storage and can be rapidly accessed. The overall algorithm takes a few seconds for 5-input functions to about a minute for 6-input functions on a single thread of an *AMD Ryzen 9 3900X* processor.

## IV. Use Cases and Computational Package

For practical applications of high-effort optimization for small circuits, the most impactful area of application is synthetic biology. Among available design-automation platforms for living systems, *Cello* has become a de facto standard for compiling digital specifications into sequence-level genetic implementations [31]. In *Cello*, a designer writes the intended behavior in a hardware description language (e.g., Verilog) or as Boolean truth tables; the compiler then performs technology mapping to a library of characterized transcriptional NOT/NOR gates for a chosen host chassis. Gate models encode transfer functions and context constraints (e.g., dynamic range, leakage, and signal compatibility), and the tool searches gate assignments and interconnections that maximize a particular scoring function. The output is a collection of promoters, ribosome binding sites, coding sequences (e.g., repressors or guide RNAs), and terminators packaged as files ready for synthesis and assembly. Because NOR/NOT families are functionally complete and well supported experimentally, *Cello* frequently realizes networks as multi-level NOR logic, aligning naturally with the architectures considered in this work [31], [35].

This workflow can prove especially useful in synthetic biology where per-cell gate budgets are tight: every additional logical gate implies extra promoters, regulatory proteins or gRNAs, and resource load, which can push designs past feasibility (e.g., with CRISPRi/dCas9 systems) [35]. Consequently, reducing the gate count at the logic level directly improves implementability, reliability, and burden. Our hybrid logic minimizer is complementary to *Cello*: it can be used as a pre-mapping pass that accepts a high-level Boolean specification (or an AIG/NIG netlist) and emits a logically equivalent NOR2 network.

Starting from a Boolean specification (behavioral Verilog) the hybrid logic minimizer module iteratively identifies eligible substructures within a larger heuristic circuit and evaluates their local Boolean expressions to compute a canonical 16-bit truth-table index. The index queries the precomputed library to retrieve the minimal realization. The algorithm then reconstructs the gate tree, applying the required input permutations and verifying the gate count. It then emits a structural Verilog netlist composed exclusively of two-input NOR primitives. The resulting NOR-only netlist is provided to Cello-v2 along with the relevant inputs (user constraint file, input-sensor, and output-device specifications).

## V. Discussion and Future Directions

Our results show that even a limited 4-input database has the potential to provide gate savings for much larger circuits and thus could potentially be a valuable addition to standard logic synthesis tools. The 4-input results shown in Fig. 3 also suggest that this is far from the maximum possible improvement. From the results that were obtained, it is clear that oftentimes a possible local gain doesn’t lead to an overall gain, which explains why there is such a high percentage of 5-input functions that could not be improved as the algorithm currently stands. The primary challenge resides in the fact that optimal circuits oftentimes have gates with large fan-out values (the number of gate inputs driven by the output of this logical gate) to contribute to an overall saving in the number of unique gates. In a typical *ABC* output, there remain a fair amount of redundancies in the calculated circuits. By locally replacing a subcircuit to make it more efficient, one can actually make things less efficient elsewhere due to the fan-out problem.

In future iterations of this work, we will explore ways to overcome this challenge of having gates with fan-outs greater than 1. When searching for subcircuits to replace with those of the database, one can factor in this fan-out number into the algorithm to add a global perspective to the local search. Also, an optimal implementation does not necessarily mean that it is the one that will lead to the greatest global gain in gate number for the circuit. Thus, an interesting direction is to look for near-optimal subcircuits that contain redundant logic with respect to the whole circuit. While in this work we only looked at one implementation per Boolean function, the *BESS* algorithm is able to provide all the alternative solutions given desired criteria. The promising results shown here also show that there is a real value to expanding the database of functions with 5 inputs or more. Having 5-input optimal circuits would lead to far more efficient simplifications but also lead to having a database that would be extremely large in size. Given that 4-input optimal circuits can nowadays be computed with relative ease, 5-input optimal implementations are within reach but it will also be unlikely that one can go much beyond that while maintaining this optimal criterion. Finally, the algorithm here looks at the extremities first and moves towards the “root” of the circuit. A more thorough search of substructure would most likely lead to better results, so there are promising directions with respect to the proposed algorithm as well.

## References

[1] R. K. Brayton, G. D. Hachtel, C. McMullen, and A. Sangiovanni-Vincentelli, Logic minimization algorithms for VLSI synthesis. Springer Science & Business Media, 1984, vol. 2.

[2] R. K. Brayton, G. D. Hachtel, and A. L. Sangiovanni-Vincentelli, “Multilevel logic synthesis,” Proceedings of the IEEE, vol. 78, no. 2, pp. 264–300, 1990.

[3] R. L. Rudell and A. Sangiovanni-Vincentelli, “Multiple-valued mini-mization for pla optimization,” IEEE Transactions on Computer-Aided Design of Integrated Circuits and Systems, vol. 6, no. 5, pp. 727–750, 1987.

[4] E. A. Ernst, “Optimal combinational multi-level logic synthesis,” Ph.D. dissertation, University of Michigan, 2009.

[5] R. L. Ashenhurst, “The decomposition of switching functions,” in Proceedings of an international symposium on the theory of switching, April 1957, 1957.

[6] H. A. Curtis, “A generalized tree circuit,” Journal of the ACM (JACM), vol. 8, no. 4, pp. 484–496, 1961.

[7] R. M. Karp, F. McFarlin, J. P. Roth, and J. Wilts, “A computer program for the synthesis of combinational switching circuits,” in 2nd Annual Symposium on Switching Circuit Theory and Logical Design (SWCT 1961). IEEE, 1961, pp. 182–194.

[8] J. P. Roth, “Minimization over boolean trees,” IBM Journal of Research and Development, vol. 4, no. 5, pp. 543–558, 1960.

[9] J. P. Roth and R. M. Karp, “Minimization over boolean graphs,” IBM journal of Research and Development, vol. 6, no. 2, pp. 227–238, 1962.

[10] L. Hellerman, “A catalog of three-variable or-invert and and-invert logical circuits,” IEEE Transactions on Electronic Computers, no. 3, pp. 198–223, 1963.

[11] R. A. Smith, “Minimal three-variable nor and nand logic circuits,” IEEE Transactions on Electronic Computers, no. 1, pp. 79–81, 1965.

[12] R. Drechsler and W. Günther, “Exact circuit synthesis,” in In Int’l Workshop on Logic Synth. Citeseer, 1998.

[13] S. Muroga and T. Ibaraki, “Design of optimal switching networks by integer programming,” IEEE Transactions on Computers, vol. 100, no. 6, pp. 573–582, 1972.

[14] E. S. Davidson, “An algorithm for nand decomposition under network constraints,” IEEE Transactions on Computers, vol. 100, no. 12, pp. 1098–1109, 1969.

[15] T. Nakagawa, H. Lai, and S. Muroga, “Design algorithm of optimal logic networks by the branch-and-bound approach.” J. COMP. AIDED VLSI DES., vol. 1, no. 2, pp. 203–231, 1989.

[16] J. A. Darringer, W. H. Joyner, C. L. Berman, and L. Trevillyan, “Logic synthesis through local transformations,” IBM Journal of Research and Development, vol. 25, no. 4, pp. 272–280, 1981.

[17] A. Vincentelli, A. Wang, R. Brayton, and R. Rudell, “Mis: Multiple level logic optimization system,” IEEE Transactions on Computer Aided Design of Integrated Circuits and Systems, 1987.

[18] K. Bartlett, W. Cohen, A. De Geus, and G. Hachtel, “Synthesis and optimization of multilevel logic under timing constraints,” IEEE Trans. Computer-Aided Design, vol. 5, no. 4, pp. 582–596, 1986.

[19] K. A. Bartlett, R. K. Brayton, G. D. Hachtel, R. M. Jacoby, C. R. Morrison, R. L. Rudell, A. Sangiovanni-Vincentelli, and A. Wang, “Multi-level logic minimization using implicit don’t cares,” IEEE Transactions on Computer-Aided Design of Integrated Circuits and Systems, vol. 7, no. 6, pp. 723–740, 1988.

[20] D. Brand, “Logic synthesis,” in Design Systems for VLSI Circuits: Logic Synthesis and Silicon Compilation. Martinus Nijhoff, 1987, pp. 301– 326.

[21] G. Hachtel, M. Lightner, K. Bartlett, D. Bostwick, R. Jacoby, P. Moceyunas, C. Morrison, X. Du, and E. Schwarz, “Bold: The boulder optimal logic design system,” in Proceedings of the Twenty-Second Annual Hawaii International Conference on System Sciences. Volume 1: Architecture Track, vol. 1. IEEE Computer Society, 1989, pp. 59– 60.

[22] R. K. Brayton, Algorithms for multi-level logic synthesis and optimization. IBM Thomas J. Watson Research Center, 1986.

[23] L. Berman and L. Trevillyan, “A global approach to circuit size reduction,” in Proceedings of the fifth MIT conference on Advanced research in VLSI, 1988, pp. 203–214.

[24] C. L. Berman and L. H. Trevillyan, “Improved logic optimization using global-flow analysis,” in The Best of ICCAD. Springer, 2003, pp. 217– 225.

[25] E. Sentovich, K. Singh, L. Lavagno, C. Moon, R. Murgai, A. Saldanha, H. Savoj, P. Stephan, R. K. Brayton, and A. L. Sangiovanni-Vincentelli, “Sis: A system for sequential circuit synthesis,” EECS Department, University of California, Berkeley, Tech. Rep. UCB/ERL M92/41, May 1992. [Online]. Available: http://www2.eecs.berkeley.edu/Pubs/TechRpts/1992/2010.html

[26] E. M. Sentovich, K. J. Singh, C. Moon, H. Savoj, R. K. Brayton, and A. Sangiovanni-Vincentelli, “Sequential circuit design using synthesis and optimization,” in Proceedings 1992 IEEE International Conference on Computer Design: VLSI in Computers & Processors. IEEE, 1992, pp. 328–333.

[27] A. Kuehlmann, V. Paruthi, F. Krohm, and M. K. Ganai, “Robust boolean reasoning for equivalence checking and functional property verification,” IEEE Transactions on Computer-Aided Design of Integrated Circuits and Systems, vol. 21, no. 12, pp. 1377–1394, 2002.

[28] R. Brayton and A. Mishchenko, “Abc: An academic industrial-strength verification tool,” in International Conference on Computer Aided Verification. Springer, 2010, pp. 24–40.

[29] A. M. R. Brayton and A. Mishchenko, “Scalable logic synthesis using a simple circuit structure,” in Proc. IWLS, vol. 6. Citeseer, 2006, pp. 15–22.

[30] J. A. Brophy and C. A. Voigt, “Principles of genetic circuit design,” Nature methods, vol. 11, no. 5, pp. 508–520, 2014.

[31] A. A. Nielsen, B. S. Der, J. Shin, P. Vaidyanathan, V. Paralanov, E. A. Strychalski, D. Ross, D. Densmore, and C. A. Voigt, “Genetic circuit design automation,” Science, vol. 352, no. 6281, 2016.

[32] M. A. Al-Radhawi, A. P. Tran, E. A. Ernst, T. Chen, C. A. Voigt, and E. D. Sontag, “Distributed implementation of boolean functions by transcriptional synthetic circuits,” ACS Synthetic Biology, vol. 9, no. 8, pp. 2172–2187, 2020, pMID: 32589837. [Online]. Available: 10.1021/acssynbio.0c00228

[33] Y. Chen, S. Zhang, E. M. Young, T. S. Jones, D. Densmore, and C. A. Voigt, “Genetic circuit design automation for yeast,” Nature Microbiology, vol. 5, no. 11, pp. 1349–1360, 2020.

[34] P.-F. Xia, H. Ling, J. L. Foo, and M. W. Chang, “Synthetic genetic circuits for programmable biological functionalities,” Biotechnology advances, vol. 37, no. 6, p. 107393, 2019.

[35] M. W. Gander, J. D. Vrana, W. E. Voje, J. M. Carothers, and E. Klavins, “Digital logic circuits in yeast with crispr-dcas9 nor gates,” Nature communications, vol. 8, no. 1, pp. 1–11, 2017.

[36] D. D. Jatkar, M. A. Al-Radhawi, C. A. Voigt, and E. D. Sontag, “Modeling and minimization of dcas9-induced competition in crispri-based genetic circuits,” bioRxiv, pp. 2025–11, 2025.

[37] S. Cho, D. Choe, E. Lee, S. C. Kim, B. Palsson, and B.-K. Cho, “High-level dcas9 expression induces abnormal cell morphology in escherichia coli,” ACS synthetic biology, vol. 7, no. 4, pp. 1085–1094, 2018.

[38] J. Shin, S. Zhang, B. S. Der, A. A. Nielsen, and C. A. Voigt, “Programming escherichia coli to function as a digital display,” Molecular systems biology, vol. 16, no. 3, p. e9401, 2020.

[39] J. P. Padmakumar, J. J. Sun, W. Cho, Y. Zhou, C. Krenz, W. Z. Han, D. Densmore, E. D. Sontag, and C. A. Voigt, “Partitioning of a 2-bit hash function across 66 communicating cells,” Nature Chemical Biology, vol. 21, no. 2, pp. 268–279, 2025.

